# Hierarchical Signatures of Language in the Human Brain

**DOI:** 10.64898/2026.02.15.705992

**Authors:** Nicole Eichert, Charlotte Favre, Donna Gift Cabalo, Alexander Ngo, Jordan DeKraker, Youngeun Hwang, Meaghan Smith, Vivian Crooks, Paul Bautin, Raul Rodriguez-Cruces, Kate E. Watkins, Oiwi Parker Jones, Casey Paquola, Saad Jbabdi, Boris C. Bernhardt

## Abstract

Language relies on a hierarchy of sensory and cognitive processes, yet how different levels of this hierarchy are supported by distinct neural architectures remains unclear. Here we show that semantic processing, compared with phonological processing, is associated with higher-level brain networks, as characterized by resting-state fMRI connectivity, *in vivo* measures of cortical myelin, and cortical types derived from a cytoarchitecture-defined reference atlas. These relationships were established using individualized ultra-high field fMRI in both English and French speakers leveraging a multi-session, multi-modal 7T MRI protocol including a language localizer. For comparison, we developed an artificial neural network, in which a representational hierarchy spontaneously emerged where phonological information was captured in earlier layers and semantic information in later layers. By integrating individualized functional mapping, neuroanatomical characterization and artificial intelligence, this study advances understanding of the neural basis of language and provides a framework for linking biological and artificial systems of communication.

**Significance Statement:** Language is widely described as hierarchical, yet how this functional organization is implemented in the brain’s biological architecture remains unclear. By combining ultra–high field, individualized neuroimaging with in vivo measures of cortical microstructure and large-scale connectivity, this study establishes a framework for linking distinct levels of linguistic computation to the brain’s structural and functional organization. Integrating these findings with artificial neural network modeling further reveals shared principles between biological and machine systems. Together, this work advances a biologically grounded account of language, bridges cognitive neuroscience and artificial intelligence, and provides a roadmap for understanding how complex cognition emerges from structured brain architecture.

## Introduction

Language is one of the most extensively studied brain functions, in both clinical (Bookheimer 2002; Saur and Hartwigsen 2012) and basic research settings (Fedorenko et al. 2011; Hickok 2012), owing to its central importance for the human experience (Tomasello 2009; Fitch 2010). It draws on a wide range of sensory and cognitive processes, and the neural basis of language therefore provides a unique window into human thought (Fodor 1979), the coupling of brain structure and function (Friederici 2011), and even the evolutionary history of the human brain (Mars et al. 2018; Eichert and Mars 2025). Therefore, research on the neurobiology of language has long been at the forefront of cognitive and neuroanatomical sciences. More recently, interest in the field has been further driven by the prospect that integrating neuroanatomy with artificial intelligence could yield *in-silico* models of the brain and shed light on its functional and computational architecture (Caucheteux et al. 2023).

Language is an umbrella term for a set of interacting cognitive processes, each with distinct developmental trajectories (Friederici et al. 2017), clinical manifestations (Dronkers et al. 2017), and neurobiological substrates (Price 2012). Often, these processes are described as following a hierarchical organization, in which successive levels build on and depend upon earlier ones (**Figure 1A**). Such a progressive sequence, spanning low-to high-level cognition, runs from sensory processes, phonetics and phonology through morphology, syntax and semantics, to pragmatics (Friederici 2012; Hagoort 2019). It remains an open question, however, to which extent such a graded progression of linguistic features maps onto hierarchical motifs observed across the cortex.

**Figure 1:**
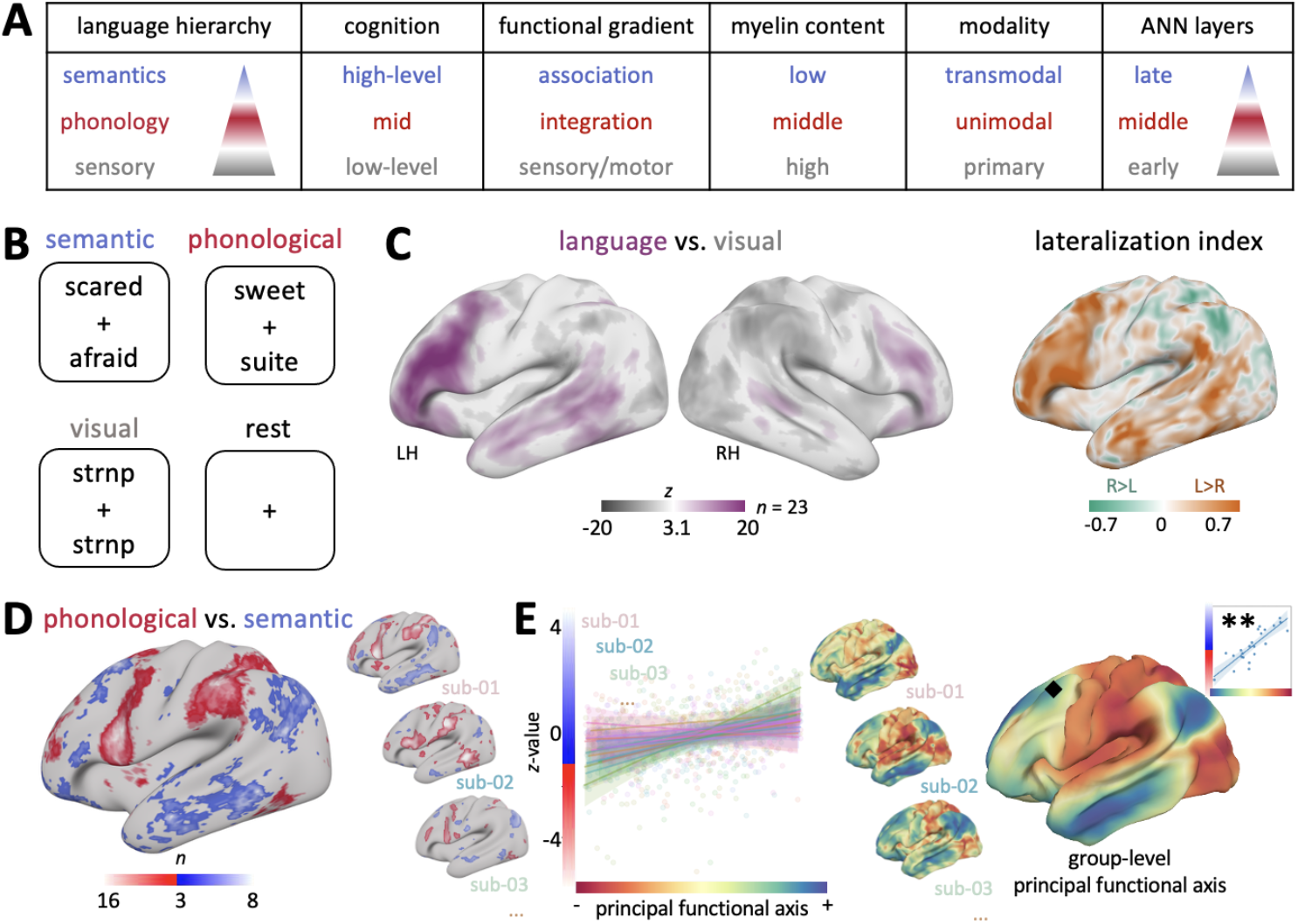
Language hierarchy. **A**: Proposed relations between levels of language hierarchy, descriptors of the brain and ANN layers. The coloured triangle represents levels of a conceptual hierarchy replicated at all descriptors. **B**: Behavioural task paradigm. For each condition, an example ‘yes’-trial is shown. **C**: *Left:* Group-level fMRI activation map for the semantic *plus* phonological *minus* visual condition (*n*=23, fixed-effects analysis, threshold: *z*=3.1). *Right:* Vertex-wise lateralization index. **D**: Count map, summing binary and thresholded statistical maps from the phonological *minus* semantic contrast (individual-subject threshold: *z*=3.1, count map threshold: *n*=3). Three example individual participant maps are shown. **E**: Correlation of individual *z*-maps and individual resting-state principal embeddings (*p*<0.01 in 18 of 23 participants, corrected for spatial auto-correlations). Three example participant maps are shown and the group-level resting-state embedding. Diamond and inset (*right*): A group-level vertex-wise permutation test demonstrates a spatial link between *z*-values and principal functional axis (*p*<0.01, *n*=23, family-wise-error correction using FSL’s randomise).

Understanding functional hierarchies, initially described in non-human animals (Rockland and Pandya 1979; Felleman and Van Essen 1991; Mesulam 1998; Hilgetag and Goulas 2020; Vezoli et al. 2021), in humans has motivated increasingly sophisticated modelling of non-invasive neuroimaging (Margulies et al. 2016; Smallwood et al. 2021; Bernhardt et al. 2022). Currently, the field is furthermore witnessing a shift from broad group-level characterizations toward precise and individualized mapping of language networks with their macro- and micro-anatomical underpinnings (Poldrack et al. 2008; Braga and Buckner 2017; Gratton et al. 2018; Eichert et al. 2021; Gordon et al. 2023). Advances in ultra-high-field magnetic resonance imaging (MRI), particularly innovations to image the living human brain at 7 Tesla (7T), have substantially improved signal, measurement fidelity, and spatial specificity. This has led to measurement gains for both structural and functional imaging, enabling more precise and biologically valid mapping of individual brain networks (Cabalo et al. 2025; Paquola et al. 2025) as well as surface-based analysis frameworks (Robinson et al. 2014; Glasser et al. 2016; Cruces et al. 2022; Larivière et al. 2023). Building on the concept of anatomically localized specialization, however, the question remains of how the hierarchical organization of the distributed language networks aligns with the brain’s large-scale functional axes (Paquola et al. 2022, 2025).

At a finer scale, the brain’s microstructural architecture provides the biological substrate that constrains and enables functional specialization. Variations in cortical microstructure, such as intracortical myelin content, laminar differentiation, and cytoarchitecture, can shape the representational capacity of cortical areas by modulating their integrative properties (Godlove et al. 2014). In recent years, these microarchitectural properties have been increasingly visualized in the living human brain using myelin-sensitive imaging contrast (Paquola and Hong 2023). Cortex-wide analyses of microstructural metrics have robustly demonstrated macroscale gradients running from sensory to transmodal areas that align with foundational descriptions on how information is processed across the cortical hierarchy (Glasser and Van Essen 2011; Margulies et al. 2016; Burt et al. 2018; Huntenburg et al. 2018; Paquola, Bethlehem, et al. 2019; Bernhardt et al. 2025). In the language domain, accordingly the functional hierarchy, from perceptual to phonological and semantic processes, emerges in part from underlying structural constraints, including myeloarchitectural differences . Connectional and myeloarchitectural differences between the hemispheres, between posterior and frontal language areas and across the dorsal/ventral language stream have been shown to be to support distinct different levels of a language hierarchy. For example, connectional and myeloarchitectural differences between the hemispheres, between posterior and frontal language regions, and across the dorsal and ventral language streams have been shown to support distinct levels of a language hierarchy (Hagoort and Indefrey 2014; Friederici 2017; Yuan et al. 2021). These observations suggest that language hierarchies are embedded within broader gradients of cortical organization, motivating a closer examination of how functional language processes relate to intrinsic microstructural and systems-level properties of the brain.

Artificial neural networks provide a useful framework to examine whether and how hierarchical principles observed in the human language system can emerge from language-focused learning alone. Shared modelling revealed that representational and computational properties of human language processing can be recapitulated in artificial models (Caucheteux et al. 2022; Goldstein et al. 2022). For example, encoding models based on sequential contextual embeddings from transformer architectures reproduce key temporal characteristics of human language processing (Goldstein et al. 2025). These findings suggest that such hierarchies may reflect general solutions to linguistic information processing rather than exclusively biological mechanisms. Here, we aim to extend these insights using representational similarity analysis (RSA), to test whether a division between phonological and semantic representations also emerges as an advantageous strategy in an artificial neural network.

In this paper, we investigated the relationships of linguistic processing, function and microstructure of the brain as well as representational hierarchies using multimodal and individualized brain mapping. The study had three main objectives. First, we sought to generate robust activation maps for phonological and semantic processing, using a 7T precision mapping fMRI paradigm. Second, we examined how language activation relates to large-scale intrinsic functional and microstructural markers of brain organization and hierarchies, testing the hypothesis that semantic processing is supported by high-level, less myelinated cortical regions compared with those implicated in phonological processing. Third, we asked whether a similar division of labour—between lower- and higher-level information—can also emerge as an advantageous strategy in an artificial neural network trained to perform the same language task.

## Results

This study integrated ultra–high-field 7T functional and microstructural MRI analysis, neuroanatomical contextualization, and artificial neural network modelling to investigate the neural architecture of language processing. Twenty-three healthy adults performed a word judgment task involving semantic and phonological decisions during fMRI acquisition, in either English (*n*=14) or French (*n*=9). Individual task activation patterns were related to individual resting-state functional gradients, quantitative T1 (qT1) maps, and a reference atlas of cortical types to characterize structure–function relationships along the cortical hierarchy. Finally, an artificial neural network was designed to perform analogous semantic and phonological classifications of word pairs, enabling comparison of representational organization between biological and artificial systems.

### Behavioural Performance

We adapted an established linguistic task paradigm that required participants to make a phonological, semantic, or visual decision about word pairs during the fMRI scan (Gough et al. 2005). Behavioural analysis demonstrated that all participants engaged in the task as indexed by overall high accuracy (**Supplementary Figure S1A**, mean: 86%, min: 74%). As expected, the visual matching condition was easiest with shortest reaction times and highest accuracy (ANOVA and post-hoc *t*-test, *p*<0.001). Accuracy in the semantic condition was lower compared with the phonological condition (85% and 0.79%, *p*<0.05), whilst reaction times were not different between the conditions (*p*=0.1). The difference between conditions was expected based on the original task description (Devlin et al. 2003), and likely reflects variability in how participants precisely judged the similarity of synonym pairs, as both “yes” and “no” responses can often be considered valid. Taken together, these results show that our adaptation of this linguistic paradigm effectively probed both semantic and phonological processing, whilst difficulty was comparable between conditions.

### fMRI Language Activations

While participants performed the linguistic task, we acquired ultra-high-resolution 7T fMRI data, which was processed using a customized surface-based pipeline (Cruces et al. 2022) and statistically analysed with a general linear model using nilearn (Abraham et al. 2014). The overall language contrast (*i*.*e*. the sum of phonological and semantic condition contrasted with the visual control condition), revealed a widespread language network with pronounced activations in the inferior frontal lobes and the middle temporal lobe (**Figure 1C - left**), which matched the expected pattern from established literature (Hickok and Poeppel 2007; Price 2012). Importantly, task activations were clearly lateralized, with strongest left-lateralization in the frontal and temporal lobe (**Figure 1C - right**), which is a hallmark of the neurobiology of language (Price 2012). The visual condition, compared with rest, preferentially activated posterior regions in the occipital and parietal lobe, which is consistent with previous studies from the reading literature (Jobard et al. 2003). Calculating the consistency of the language activation map demonstrated that the two task runs resulted in highly similar activation maps within subjects, and to a significantly lower extent across subjects reflecting idiosyncratic differences (**Supplementary Figure S1C**, Spearman *r*=0.40 and 0.17, *p*<0.001). Activation maps between the English and French participants were comparable considering within-language and across-languages activation-map correlations (**Supplementary Figure S1D**, Spearman *r*=0.23 and 0.21, *p*>0.1). Our fMRI data demonstrate that our linguistic task adaptation serves as a valid localizer for cortical language areas: it results in robust activation maps within participants but is also able to pick up idiosyncratic differences across participants.

The phonological *vs*. semantic contrast revealed two distinct and distributed networks for the two task conditions (**Figure 1D, Supplementary Figure S1B**). Task activation maps are displayed as count maps, *i*.*e*. summed maps of individual binary activation maps statistically thresholded at *z*=3.1, to allow for a quantification of inter-individual variability. The phonological network consistently activated the posterior part of the inferior frontal gyrus (pars opercularis), corresponding to Brodmann Area (BA) 44, and the angular gyrus (BA 37). Further notable activations in the phonological condition were in the anterior part of the middle frontal gyrus (BA 46) and supplementary motor cortex on the medial surface. The semantic condition overall resulted in lower count values indicating more variability. Notable activations that are clearly visible on the group-level were in inferior frontal gyrus (pars triangularis, corresponding to BA 45), a more anterior ventral location (pars orbitalis) corresponding to BA 47, supramarginal gyrus, anterior temporal pole and at least one activation cluster in the middle temporal gyrus. These language-related brain activations were expected based on the literature (Hickok and Poeppel 2007; Price 2012). Indeed, we could identify robust associations of the overall language activation map to *Neurosynth* derived meta-activation maps (Yarkoni et al. 2011) related to the terms ‘language’, ‘semantics’, ‘phonological’ (**Supplementary Figure S1E**), and identified specific enrichment for the term ‘phonological’ terms when decoding the phonological *vs*. semantic contrast and for the term ‘semantics’ when decoding its inverse.

### Functional and microstructural markers

For each participant, we related the spatial pattern of language activations to functional and microstructural features of the brain, thereby characterizing how language is reflected in the brain’s broader architecture. First, we assessed how the statistical task map from the phonological *vs*. semantic contrast related to the spatial map of the principal functional axis. We observed a strong association at the group level (*r*=0.54, *p*<0.01, controlled for spatial auto-correlation using spin tests (Alexander-Bloch et al. 2018) implemented in BrainStat, Larivière et al. 2023) and also using each individual’s task and gradient score maps (**Figure 1E**): A significant positive correlation was found in 18 out of 23 participants (average *r*=0.30, *p*<0.01, corrected). This result indicates that the phonological network co-localized with the apex of the gradient associated with more low-level features and local signal processing (Margulies et al. 2016). Respectively, semantic processing is associated with the apex of the gradient related to long-range connections and high-level cognition (Mesulam 1998). In addition to the individualized analysis, we also tested in what parts of the brain the strength of the task activation, *i*.*e*. fMRI *z*-value, and resting-state gradient score correlated across subjects. We found a significant association between task activation and resting-state gradient score bilaterally in the middle frontal gyrus [diamond highlight and right inset in **Figure 1E**, *p*<0.01, corrected for family-wise error using FSL’s randomise (Winkler et al. 2014)], further demonstrating that language activation is coupled to and supported by the brain’s state during rest.

Next, we assessed the relationship to qT1, which serves as an MRI-derived microstructural marker associated with overall cortical myelin content (Turner 2019), a cortical feature that is generally associated with more early sensory and perceptual processing (Glasser and Van Essen 2011; Lutti et al. 2014; Paquola et al. 2020, 2025; Sydnor et al. 2021). We also probed the relationship to the principal spatial axis of the qT1 signal profiles across the cortical ribbon, which maps a system-level gradient of microstructural differentiation ranging from sensory to transmodal areas (Paquola, Vos De Wael, et al. 2019; Cabalo et al. 2025). We found that phonological processing is localized in parts of the cortex that have lower mean qT1 values, indicative of overall higher cortical myelin content (**Figure 2B - left**, *p*<0.05). Semantic processing, respectively, localizes in cortical areas of lower cortical myelin content, overlapping with the high-level apex of the qT1 gradient. We observed a strong association of the strength of task activation, *i*.*e*., the *z*-value, with myelin content quantified on within both the phonological and the semantic network (**Figure 2C - left**): Lower qT1 values, *i*.*e*. higher cortical myelin content, is associated with higher statistical task activation (mixed–effects model, Cohen’s *d*=–0.43, *p*<0.01). Statistical task maps and individual qT1 maps were not correlated on a vertex-wise level, however, (average *r*=0.05), suggesting that the relationship mainly exists on the global network, rather than on the local level.

**Figure 2:**
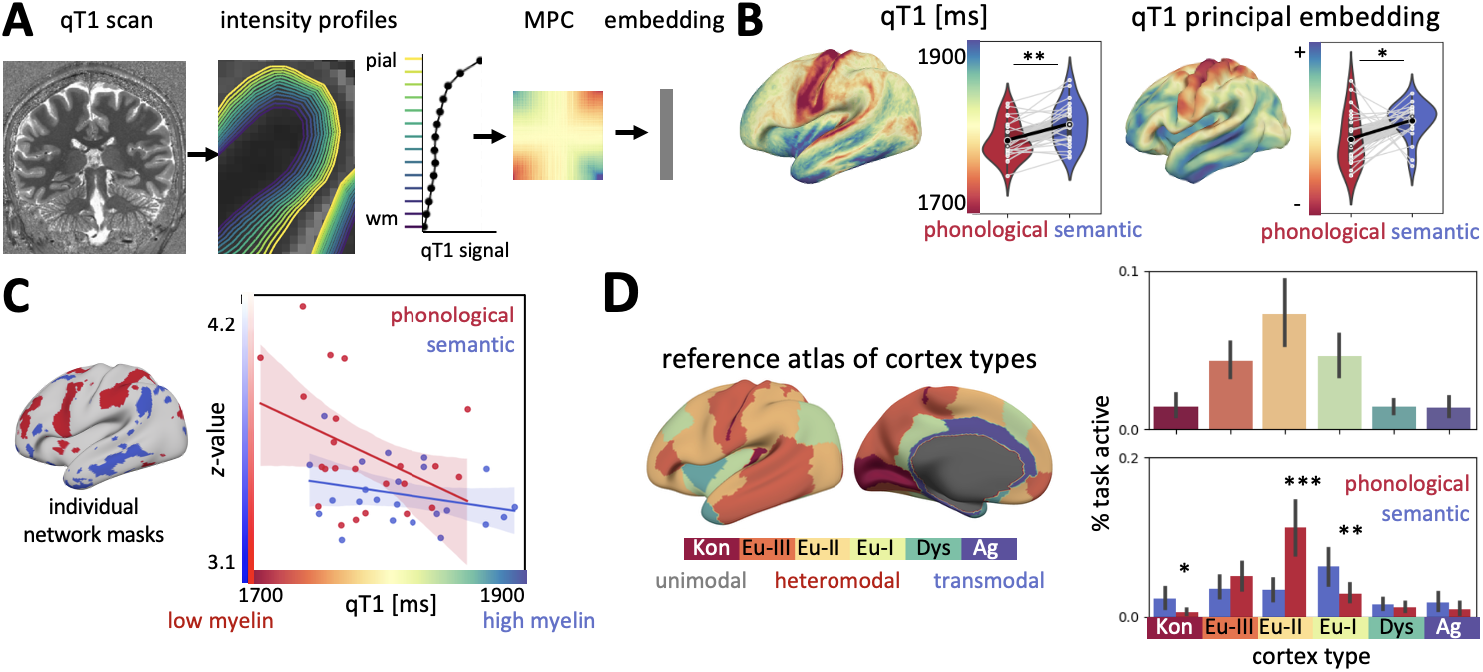
Levels of language hierarchy are supported by distinct subsystems. **A**: Quantitative T1 relaxometry (qT1) profiling, and cortical profiles and a brain-wide gradient. **B**: Phonological and semantic task-activation localize in cortical areas differing in mean qT1 and gradient score. **C**: Association between qT1 and *z*-values. **D**: Phonological and semantic task-activation co-localize in brain areas of different cortical types.

We confirmed and extended these findings by integrating a *post-mortem* reference atlas of cortical types (García-Cabezas et al. 2020) (**Figure 2D**). The proportion of task activation within a segmentation of cortical types revealed a double-dissociation: Semantic processing preferentially activating areas of type-1 eulaminate cortex, which is characterized by a more well-developed layer IV typically associated with higher-order processing, whilst phonological processing engaged more type-2 eulaminate areas, which have thicker granular layer IV and larger pyramidal neurons in supragranular layers. Taken together, these microstructural analyses demonstrate that semantic processing, compared with phonological processing, takes place in transmodal high-level areas of the cortex, characterized by lower myelin.

### Artificial neural network analogy

In a final analysis we tested whether the hierarchical nature of linguistic features studied above, *i*.*e*., levels of sensory, phonological, semantic processing, also spontaneously arises in an artificial system. By doing so, this analysis probes whether a separation of representations along sensory, phonological, and semantic dimensions constitutes a computationally sufficient solution for the task, even in the absence of explicit biological constraints. We designed an artificial neural network (ANN) to perform the same linguistic task as the participants, taking a word pair as input and predicting the semantic and phonological similarity of the pair. The model was implemented in PyTorch and trained on a large set of word pairs including uniformly sampled pairs ranging from low to high similarity and lists of homophones and synonyms.

The ANN achieved a binary classification accuracy of 0.85 and predicted semantic and phonological similarities that closely matched the true similarities (Spearman’s *r*=0.6; **Figure 3B**), indicating successful learning of the linguistic task. To characterize the network’s internal representations, we extracted activation patterns from each hidden layer for every word pair and quantified the representational similarity of the two words. Comparing these layer-specific similarity structures to the true pairwise similarities enabled a representational similarity analysis (RSA). This analysis revealed a hierarchical organization across layers (**Figure 3D**), with early layers primarily capturing visual information related to letter form, consistent with the character-indexed input to the model. Phonological representations remained high in intermediate layers; and semantic representations appeared only in the final layers. This progression of extracted information mirrors the hierarchy observed in the brain, with phonological processing supported by cortical regions lower on the cortical hierarchy and semantic processing supported by transmodal, or high-level areas (**Figure 3E**).

**Figure 3:**
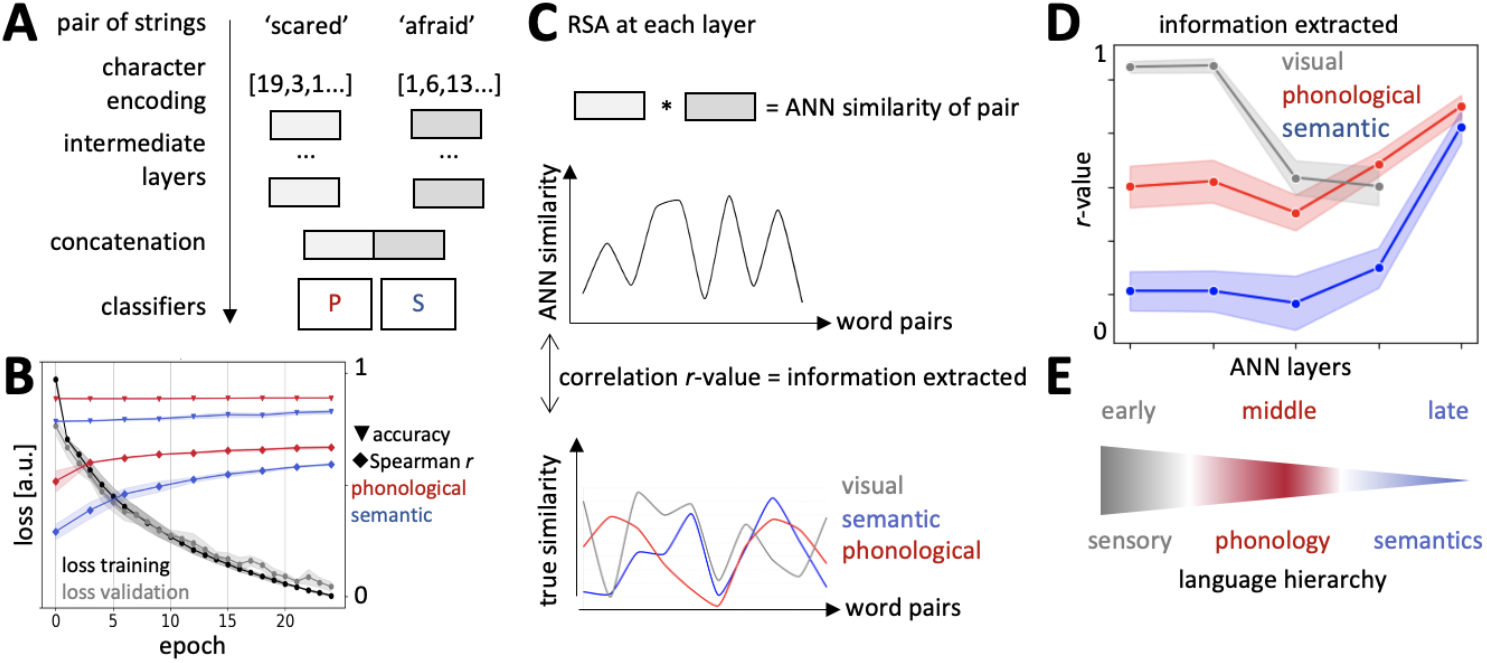
Hierarchy of information extracted in a linguistic ANN. **A**: Model architecture illustrated for one example word pair. **B**: Model performance. **C**: Representational similarity analysis (RSA) schematic to compute which type of linguistic information is processed by a given ANN layer. **D**: Types of information extracted at each ANN layer. **E**: Schematic correspondence of the levels of ANN layers and levels of the language hierarchy.

## Discussion

This study bridges cognitive neuroscience, neuroanatomy, and artificial intelligence to demonstrate that the principles of the language hierarchy, specifically the functional differentiation between lower-level phonology and higher-level semantics, is grounded in the functional and microstructural hardware of the brain. Further, we showed that such hierarchical linguistic levels can also emerge as internal representations of a simple and straightforward artificial neural network model. We established these relationships on an individualized neuroanatomical level enhancing our mechanistic insight into how biological constraints shape functional specialization in the human brain

We successfully adapted a previously established localizer task (Gough et al. 2005) for our multi-session 7T MRI protocol (Cabalo et al. 2025) to map the language system in the brain. Activations were highly robust across scanning sessions and resulted in comparable brain maps in English and French participants. Yet we observed substantial inter-subject variability, demonstrating that we were able to image idiosyncratic differences across participants at a high spatial resolution. Our results highlight the benefit of using ultra high-field 7T MRI as suitable means to localize functional brain systems at high precision (Uğurbil et al. 2013; Uğurbil 2021; Gratton and Braga 2025). We observed a left-lateralized language system and distinct networks for semantic and phonological processing. These activations overlapped with those established in a term-based meta-analysis (Gorgolewski et al. 2015) confirming that we localized key brain areas for language processing (Price 2012).

We acquired additional extensive multi-modal MRI data in all participants, allowing us to study the architectural characteristics underlying semantic and phonological processing. By using a multi-modal approach, we demonstrated that the language hierarchy is reflected by local and global descriptors of brain organisation, thereby improving our mechanistic understanding of language function. We found that, on an individual-subject level, language function closely aligns with the primary resting-state gradient. Resting-state gradients represent low-dimensional spatial embeddings of functional brain organisation that capture a global transition across the cortex from sensory to association areas (Margulies et al. 2016). The primary gradient has previously been linked to a wide range of conceptual, computational and microstructural hierarchies, such as from externally to internally oriented networks (Huntenburg et al. 2018; Smallwood et al. 2021), levels of microcircuitry heterogeneity (Demirtaş et al. 2019) and progression from localized, modality-specific processing towards integrative, transmodal cognition (Mesulam 1998; Wang et al. 2025). We found that semantic processing engaged regions near the gradient apex typically linked to higher-order cognition, long-range connections, and transmodal integration (Buckner and DiNicola 2019; Ito et al. 2020; Sydnor et al. 2023; Wang et al. 2023), whereas phonological processing was associated with the opposite apex, associated with lower-order cognitive functions, short-range connectivity, and unimodal processing.

Furthermore, we found differences across conditions in mean qT1 and qT1 cortical profile embeddings. Whilst not being a direct quantitative measure of myelin, qT1 and similar proxies are sensitive to cortical myelin content (Lutti et al. 2014). We found that semantic processing takes place in brain areas containing less cortical myelin, compared to those activating during phonological processing. This finding was confirmed and extended by integrating a *postmortem* standard atlas of cortical types (von Economo 2009; García-Cabezas et al. 2020). A double dissociation emerged, with phonological processing predominantly engaging eulaminate-I regions and semantic processing preferentially activating eulaminate-II regions. On a local level, cortical myelin content inversely correlates with cell density (Paquola, Vos De Wael, et al. 2019; Chang et al. 2022), size of dendritic trees and complexity of intracortical circuitry (Patel et al. 2019; Timmler and Simons 2019). On a global level, the cortical gradient ranging from low to high myelin content captures different levels laminar elaboration (Rockland and Pandya 1979; Barbas 1986; García-Cabezas et al. 2020) and more broadly a sensory-association hierarchy (Glasser and Van Essen 2011; Nieuwenhuys 2013).

These microstructural patterns indicate that language functions align with distinct levels of the cortical hierarchy: At the lower level, which was not directly probed by our task, highly myelinated primary sensorimotor areas support perceptual aspects of language processing and downstream motor signalling for speech production. Phonological processing occupies parts of the cortex characterized by mid-to-high myelin content and large pyramidal neurons in supragranular layers, indicating a cortical makeup that supports unimodal feed-forward processing of auditory information (Rauschecker and Scott 2009) and information related to planning of speech movements (Tremblay and Deschamps 2020). Semantic decisions, at the peak of the language hierarchy, benefit from a cortical makeup that supports integration of multimodal representations via long-range connections. Indeed, multimodal conceptual representations are distributed across the cortex (Huth et al. 2012) and flexibly change based on task demands and contextual or behavioural relevance (Huth et al. 2016).

In a final analysis, we implemented an artificial neural network (ANN) to perform an equivalent semantic and phonological classification task. Input words underwent sequential processing in several ANN layers, each producing an abstract embedding representing both phonological and semantic features. Using representational similarity analysis, we found that information processing follows the same stages as expected based on the cortical language hierarchy (**Figure 1A**), thus providing us with additional mechanistic insight: Early layers of the ANN correspond to low-level sensorimotor brain areas relying on localized processing via short-range connections. These representations closely mirror the concrete stimulus features. Intermediate layers increasingly abstract the raw inputs generating unimodal representations. Later layers are analogous to high-level cortical areas that integrate pre-processed information, engaging in more distributed processing via long-range connections. We explicitly modelled symbolic linguistic features from classical psycholinguistics in a task-matched custom model, rather than using large pretrained language models whose internal representations are shaped by unconstrained text prediction objectives (Schrimpf et al. 2021; Goldstein et al. 2022; Caucheteux et al. 2023). In doing so, our framework combines individualized neuroanatomy with interpretable computational modelling, yielding a mechanistic account of how linguistic computations map onto cortical hierarchies.

## Methods

### Language task paradigm

We adapted a previous language localizer task (Devlin et al. 2003; Gough et al. 2005) for our study. Briefly, participants were required to make a judgement about a pair of written words presented simultaneously on a screen (**Figure 1B**). In the semantic condition, participants judged whether the two words were synonyms (*i*.*e*., shared the same meaning), in the phonological condition, whether they were homophones (*i*.*e*., sounded the same) and, in the visual matching condition, whether two meaningless letter strings were identical. In each trial, a fixation cross was presented for 500 ms, followed by a pair of words displayed above and below the fixation cross for 500 ms. Subsequently, a blank screen was shown for 2 seconds, during which participants indicated a yes/no response via button press with the right index or middle finger. The task followed a block-design with 10 trials per condition, and three interleaved rest-blocks, presented in a pseudo-random order, controlling for order effects (p-s-v-r-v-p-s-r-s-v-p-r; p=phonological, s=semantic, v=visual, r=rest). Each block began with a 3-seconds written instruction indicating the required judgement. The task scan lasted for 6 minutes and was repeated once in the same session using different stimuli (*i*.*e*. two runs). We used the original stimulus materials for English-speaking participants and generated a new stimulus set for French-speaking participants, with word frequency matched across the phonological and semantic conditions. All participants viewed the same stimulus set, according to their first language. Behavioural performance was evaluated to verify task engagement.

### Participants

MRI data were acquired at the McConnell Brain Imaging Centre of the Montreal Neurological Institute (MNI, The Neuro) for 23 healthy adults (13/10 males/females, age = 28.0 ± 4.58, 14/9 native English-/French-speaker), with no signs of neurological or psychiatric illness. The MRI data acquisition protocols were approved by the Research Ethics Board of McGill University and the MNI (2023-8971/2022-8526). All participants provided written informed consent, which included a provision for openly sharing all data in anonymized form.

### MRI data acquisition and pre-processing

MRI data were acquired on a 7T Magnetom Terra Siemens scanner and formed part of the openly-accessible 7T precision neuroimaging (PNI) dataset (Cabalo et al. 2025). MRI data acquisition, protocols and pre-processing have been extensively described previously (Cabalo et al. 2025). In brief, participants underwent a comprehensive multimodal scanning protocol across three different sessions, each lasting ∼90 minutes. A synthetic T1w image and a quantitative T1 relaxometry map (qT1) were both derived from a MP2RAGE sequence (0.5 mm isovoxels, matrix=320×320, 320 sagittal slices, TR=5170 ms, TE=2.44 ms, TI1=1000 ms, TI2=3200 ms, flip=4°, iPAT=3, partial Fourier=6/8, FOV=260×260 mm^2^). Two fMRI scans were acquired for the two task runs and during a 6-minutes resting-state scan. The BOLD fMRI was acquired using a 2D BOLD multi-echo echo-planar sequence (University of Minnesota, CMRR; 1.9 mm isovoxels, 75 slices oriented to AC-PC-39 degrees, TR=1690 ms, TE = 10.80/27.3/43.8 ms, flip=67°, FOV=224×224 mm^2^, slice thickness=1.9 mm, MB=3, echo spacing=0.53 ms).

### MRI data pre-processing

MRI data pre-processing was performed using *micapipe v*.*0*.*2*.*3* (http://micapipe.readthedos.io, Cruces et al. 2022), an integrated pipeline combining standard tools such as FSL (Jenkinson et al. 2012), HCP’s workbench command (Marcus et al. 2011) and ANTs (Avants et al. 2011). Following standard pre-processing of the MP2RAGE sequence, the cortical surface was extracted using FastSurfer *v*.*2*.*0*.*0* (Henschel et al. 2020) providing a standard fsLR-32k mesh. Each individual’s surface reconstruction was visually inspected for quality control and corrected. Post correction, 14 equivolumetric surfaces between pial and white matter boundaries were constructed (**Figure 2A**). Each individual’s qT1 scan intensities were sampled along these surfaces resulting in surface-based qT1 profiles. Based on these profiles, microstructural profile covariance (MPC) matrices were computed, which underwent dimensionality reduction using the Brainspace gradient function (Vos de Wael et al. 2020),using a sparsity setting of top 10% and diffusion *α* = 0.5, resulting in a qT1 principal axis, or gradient, for each individual.

FMRI data were pre-processed using TEDANA (DuPre et al. 2021) to combine multi-echo information and further standard steps, such as motion-correction, high-pass filtering, surface sampling, global signal regression and smoothing with a 10 mm Gaussian kernel prior to generating task activation maps. Vertex-wise individual functional connectome (FC) matrices were derived from each participant’s resting-state fMRI data following downsampling to a 5k-fsLR surface space (Wang et al. 2025). Each individual’s FC gradient, or principal functional axis, was derived based on the FC matrix, similar as described for the MPC matrices above, using the Brainspace gradient function. Individual-subject FC gradients were then aligned to a group-level template using Procrustes rotations to ensure consistency in ordering and polarity (Langs et al. 2015).

### fMRI task analysis

Brain activations during the language task were analysed using Python’s *nilearn* package (https://doi.org/10.5281/zenodo.8397156; (Abraham et al. 2014). A general linear model was fitted to each participant’s pre-processed surface-based fMRI data, following a smoothing step with a kernel of sigma 3 mm. The design matrix had one column for each task condition plus temporal derivatives, drifts and individual motion regressors and was convolved with a double-gamma HRF. We used a fixed-effects statistical approach, modelling each participant’s cross-session effect as first level and the group-level effect as second level. To obtain an activation map for the language system, a contrast of *phonological + semantic - visual* was obtained. A brain map for language lateralization was derived using the index (L-R)/(L+R) based on corresponding vertices from the two hemispheres, resulting in a score between −1 (maximally right lateralized) and +1 (maximally left lateralized, Bishop 2014).

Robustness of the task activations was quantified by computing the intra-subject consistency across the two task runs, using Spearman’s rank correlation coefficient, and comparing this to the inter-subject correlations. Similarly, the within- and across-group consistency for between English and French speakers was statistically compared. To obtain the semantic and phonological network, a contrast between the two conditions was computed, *i*.*e*. semantic minus phonological. A group-level count map was derived by summing all individual’s binary activation masks, thresholded at a statistical z-value of 3.1, equivalent to *p*<0.001 (Barch et al. 2013; Eichert et al. 2020).

We validated the functional relevance of our task activations using a term-based meta-analysis in *Neurosynth* (Yarkoni et al. 2011). Volumetric statistical maps from an association test, FDR-corrected at threshold 0.01, were directly downloaded from the online platform and then mapped to a standard surface. Specifically, we accessed the large-scale fMRI association test maps from the platform using the terms ‘language’, ‘phonological’ and ‘semantic’. We then computed the Dice coefficient of this map with the thresholded fixed-effects group-level activation maps from our fMRI task analysis, thresholded at *z*=3.1.

### Functional and microstructural links

The relationship between each individual’s unthresholded task activation map and their respective resting-state functional gradient was assessed using a linear regression model. We corrected for spatial auto-correlation by performing spin-tests (1000 permutations; (Alexander-Bloch et al. 2018) as implemented in BrainSpace (Vos de Wael et al. 2020). To further probe the relationship between the two metrics, we computed a vertex-wise correlation of task-based activation and resting-state gradient using FSL’s randomise (Winkler et al. 2014) providing permutation-based corrected *p*-values.

To assess associations of the language network to microstructure, we compared the mean qT1 value and the principal qT1 gradient in the semantic and phonological network across participants in a paired *t*-test. Additionally, a reference atlas of cytoarchitectural cortical types (von Economo 2009; García-Cabezas et al. 2020) was accessed via an open data repository (https://fz-juelich.sciebo.de/s/nfmYSsHjzEReYui; Paquola et al. 2025). We computed the proportion of vertices activated during semantic and phonological processing depending on each cortical type from the reference atlas of cortical types.

### Neural Network Architecture and Training

We implemented a neural network in *PyTorch* (Paszke et al. 2019) to perform the same phonological and semantic classification tasks used in the human experiments (**Figure 3A**). The model received two letter strings as input, each represented with a 28-character index encoding (26 letters plus unknown and padding). Both input vectors passed through identical processing steps across a series of layers: a 16-dimensional character embedding, long short-term memory (LSTM) encoding, two linear transformations with ReLU activation, averaging embeddings for each word across characters, followed by a linear projection. The resulting representations for the two words were then concatenated and passed to two parallel classifiers—one semantic and one phonological—each consisting of a linear layer and a sigmoid output.

For model training, we constructed a large set of English word pairs from the WordNet database (Miller 1995). Each pair was labelled based on its phonological similarity, computed using the Levenshtein distance (Levenshtein 1966), and its semantic similarity, computed with pre-trained GloVe Twitter embeddings (Pennington et al. 2014) accessed via the Python Gensim package (Řehůřek and Sojka 2010). To keep the architecture and dataset size manageable, the model was not trained to perform the visual matching task condition. Visual similarity for the representational similarity analysis (RSA) below was computed based on the number of differing characters.

The training set comprised: (1) a list of approximately 2000 homophones (accessed via https://fivejs.com/, with permission); (2) an equal number of synonyms, defined as pairs with semantic similarity > 0.9; (3) visually similar but non-homophonic words, defined as phonological similarity < 0.9; and (4) random pairings of fMRI task stimuli with other WordNet words, excluding the original task pairs. For the random pairs, we applied uniform sampling across the full range of phonological and semantic similarity from 0 to 1 (20 bins with each 500 pairs), so that each word was paired with a median of 10 other words. The dataset was augmented by adding each pair in reversed order (word1 + word2 and word2 + word1), yielding a final training set of ∼56.000 pairs. The model was trained for 20 epochs with a batch size of 32 samples and 3-fold cross-validation, using mean squared error loss and Adam optimizer. Performance was evaluated using Spearman *r* for continuous similarity metrics as well as binary classification metrics at a threshold of 0.8.

### Representational Similarity Analysis of Hidden Layers

To probe the model’s internal hierarchical organisation, we performed a representational similarity analysis (RSA) to determine which type of information is extracted at each of the network’s layers (**Figure 3C**). We presented the model 50 batches of 120 random word pairs (30 pairs each with either high or low phonological and semantic similarity) part of the training data and recorded the layer activations for each pair. At every ANN layer, we computed the cosine similarity between the two words’ internal tensor representations, yielding a similarity vector of 120 values. We then quantified how well this vector reflected true phonological, semantic, or visual similarity by calculating the Pearson correlation, producing a scalar measure of representational similarity for each layer and the three linguistic concepts.

## Supplementary Materials

**Supplementary Figure S1:**
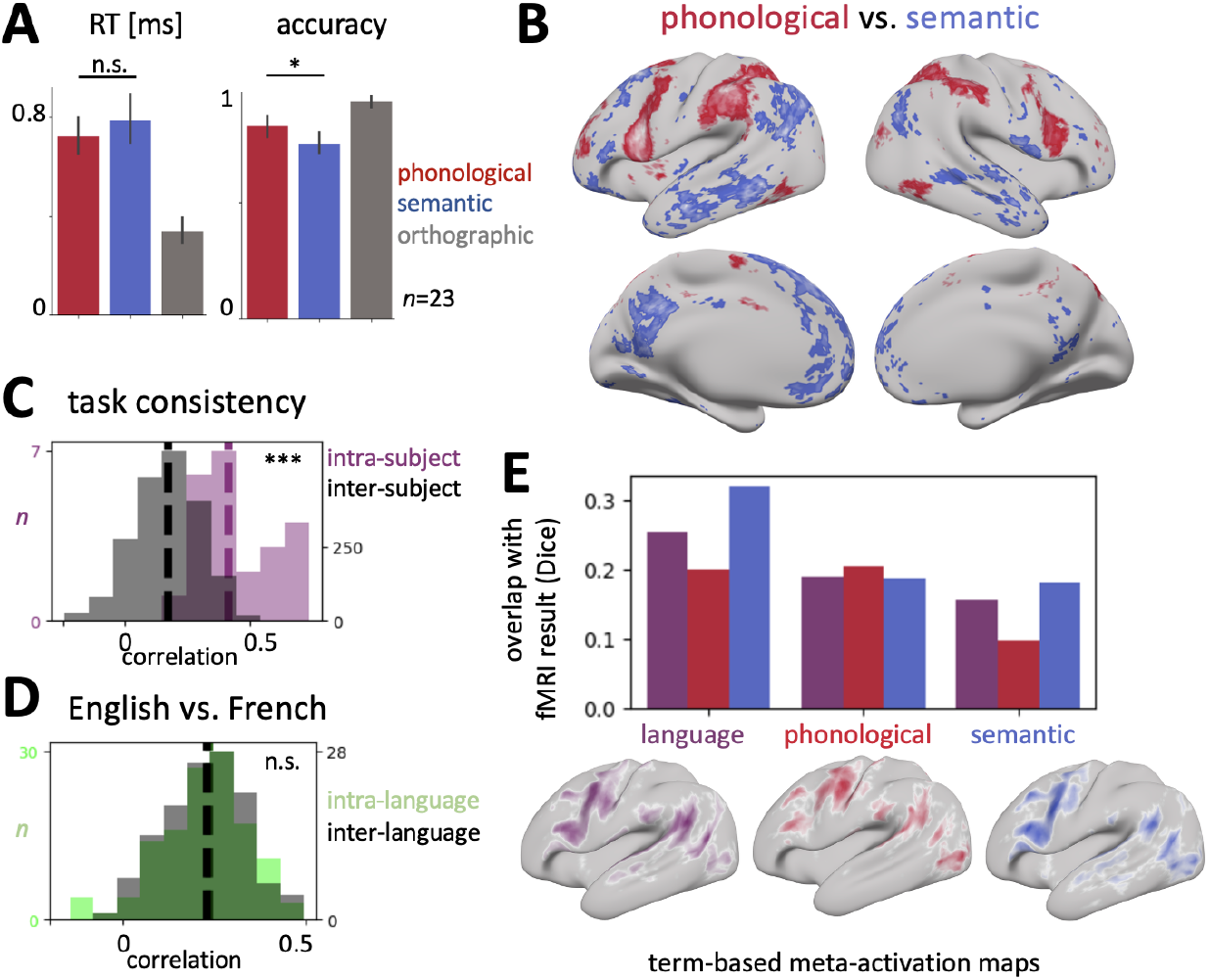
**A**: Behavioural results of task paradigm. Significance highlights for the visual condition were omitted for clarity. **B**: fMRI task count maps (extended version of main Figure 1D). **C**: The language-localizer provides comparable statistical maps across task runs within subjects (dark grey) but is able to pick up idiosyncratic differences across subjects (purple) (*p*<0.001). **D**: Task maps across English and French participants are comparable (*p*>0.5). **E**: Overlap of group-level fMRI maps and *Neurosynth* meta-activation maps.

## Acknowledgements

NE was supported by a Sir Henry Wellcome Postdoctoral Fellowship from the Wellcome Trust [222799/Z/21/Z]. JdK is supported by a Natural Sciences and Engineering Research Council of Canada - Postdoctoral Fellowship (NSERC-PDF). SJ was supported by a Wellcome Collaborative Award [215573/Z/19/Z] and a Wellcome Senior Research Fellowship [221933/Z/20/Z]. OPJ is supported by the MRC (MR/X00757X/1), Royal Society (RG\R1\241267), and ARIA (SCNI-SE01-P004). CP is supported by the Deutsche Forschungsgemeinschaft (Emmy Noether Programme – 524408221). BCB acknowledges research support from the National Science and Engineering Research Council of Canada (NSERC Discovery-1304413), CIHR (FDN-154298, PJT-174995), SickKids Foundation (NI17-039), Helmholtz International BigBrain Analytics and Learning Laboratory (HIBALL), Healthy Brains and Healthy Lives (HBHL), BrainCanada, and the Tier-2 Canada Research Chairs program. The Oxford University Centre for Integrative Neuroimaging (OxCIN) was supported by Wellcome Trust funding between 2017-2025 [203139/Z/16/Z and 203139/A/16/Z], during which time it was known as WIN (Wellcome Centre for Integrative Neuroimaging). The authors thank John F. Troutman and Joy A. Miller for providing the list of homophones used in this project. For the purpose of Open Access, the author has applied a CC BY public copyright licence to any Author Accepted Manuscript (AAM) version arising from this submission.

## Data availability

The 7T precision neuroimaging dataset is openly available via OSF (project ID: 10.17605/OSF.IO/MHQ3F) (Cabalo et al. 2025). The reference atlas of cytoarchitectural cortical types (von Economo 2009; García-Cabezas et al. 2020) can be accessed via an open data repository (https://fz-juelich.sciebo.de/s/nfmYSsHjzEReYui, Paquola et al. 2025).

## Code availability

All code generated for this project to handle the cited software tools will be deposited at https://git.fmrib.ox.ac.uk/neichert/project_semphon.

## Author Contributions

Conceptualisation: N.E. and B.C.B.; Methodology: N.E., D.G.C., J.D.K., Y.H., M.S., V.C., R R.C. and B.C.B.; Resources: D.G.C., A.G., M.S., B.C.B.; Writing—original draft: N.E. and B.C.B.; Writing— review & editing: all authors.

